# On-line reoptimization of mammalian fed-batch culture using a nonlinear model predictive controller

**DOI:** 10.1101/2022.12.28.522066

**Authors:** Tomoki Ohkubo, Yuichi Sakumura, Katsuyuki Kunida

## Abstract

Fed-batch culture enables high productivity by maintaining low substrate concentrations in the early stage of the culture to suppress the accumulation of by-products that are harmful to cell growth. Therefore, they are widely used in the production of biopharmaceuticals by mammalian cells. However, there exists a trade-off in the design of the fed-batch process: early feeding results in the accumulation of harmful by-products, whereas late feeding results in a shortage of substrates needed for cell growth and synthesis of the desired product. To manage this trade-off and maximize the product yield, model-based optimization of the feeding trajectory has been reported in several studies. A significant drawback of this off-line optimization approach is the mismatch between the predictions made using the model and the actual process states, called the process-model mismatch (PMM). If the PMM is large, the off-line optimized feeding trajectory is no longer optimal for the actual process, resulting in lower product yields. Mammalian cell culture models typically contain dozens of unknown parameters that must be estimated prior to optimization. Sufficient parameter estimation is often unachievable owing to the nonlinear nature of these models. We believe that reoptimizing the feeding trajectory in real time using a nonlinear model predictive controller (NLMPC) is an effective solution to this PMM. Although NLMPC is a model-based feedback controller widely utilised in mammalian fed-batch culture, only a few studies have applied it to on-line reoptimization, and it remains unclear whether NLMPC with a standard kinetic model can effectively compensate for a large PMM. In this study, we demonstrated the reoptimization of the feeding trajectory with a NLMPC using two previously reported standard monoclonal antibody (mAb) production models. In both models, NLMPC successfully suppressed the reduction in mAb yield caused by the intentional introduction of PMM.

## Introduction

Fed-batch culture maintains a low concentration of substrates such as glucose and glutamine in the early stage of culture to suppress the accumulation of by-products, such as lactate and ammonia, that are harmful to cells [1]. Fed-batch culture is superior to batch culture in terms of productivity with a limited amount of substrates and is widely used in the production of biopharmaceuticals.

However, there is a trade-off in the design of the feeding trajectory: feeding too early increases the accumulation of harmful by-products, whereas feeding too late causes a shortage of substrates needed for cell growth and synthesis of desired products. To manage this trade-off and maximize product yield, many studies have established mathematical models and reported their application in the optimization of feeding trajectories in the last few decades [2–9]. In these studies, the controller consisted of a model and an optimizer (Fig. 1A). The controller received the initial state of the process and predicted the state trajectory of the process using the model. Based on this prediction, the optimizer modifies the feeding trajectory such that the objective function decreases. By iterating this cycle of prediction and modification, the controller solves the optimal control problem before the actual process begins. We refer to this approach as the off-line optimization. In off-line optimization, there is no feedback from the ongoing process.

**Figure 1.**
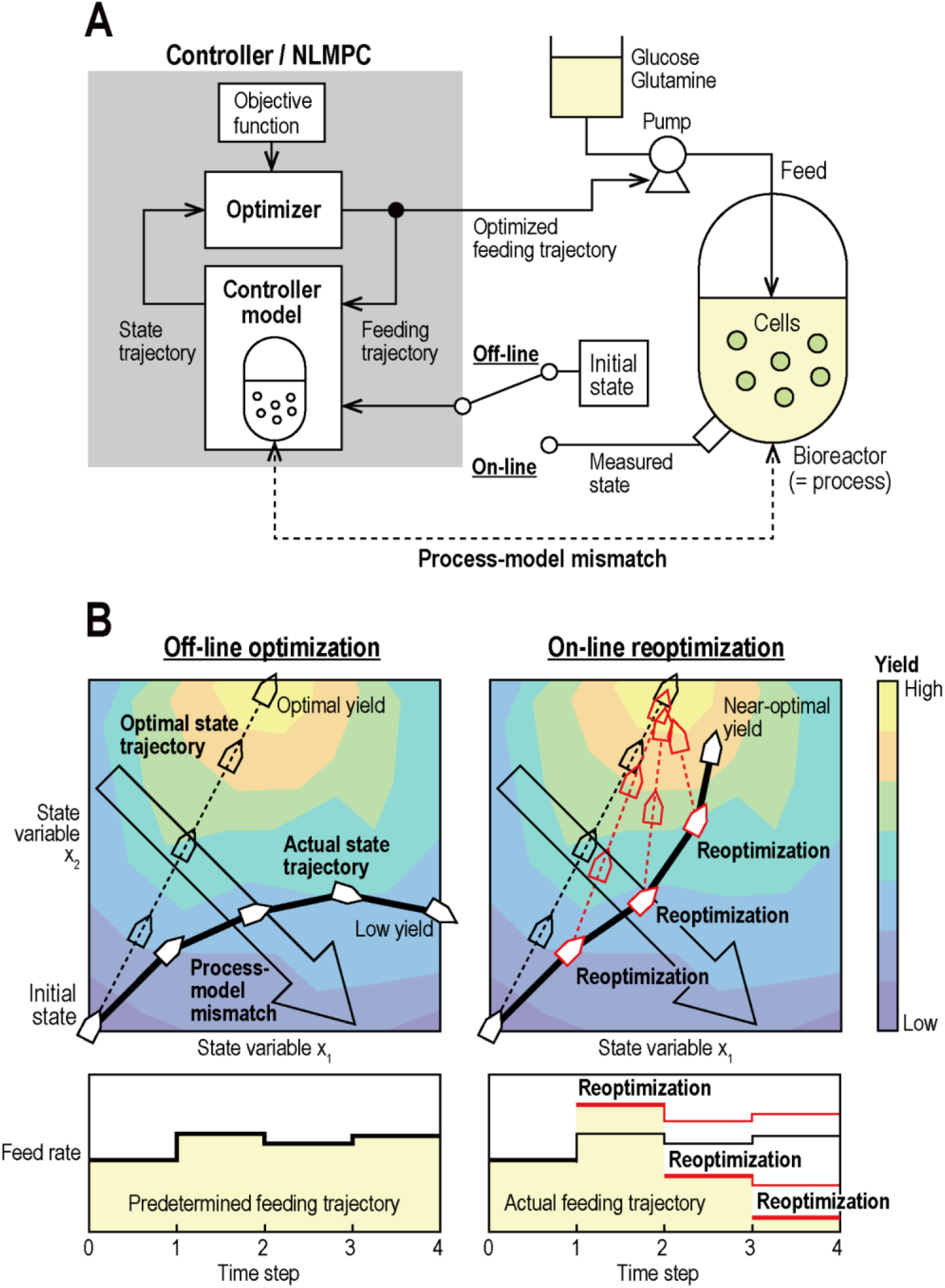
Schematic diagram of off-line optimization and on-line reoptimization. (A) The controller or NLMPC solves the optimal control problem by iterating the cycle of the prediction of the state trajectory with the model and modifying the feeding trajectory with the optimizer. In off-line optimization, the controller optimizes the feeding trajectory based on specified initial conditions before the actual process starts. In on-line reoptimization, the NLMPC receives measurements of the process state and iteratively reoptimizes the feeding trajectory based on these measurements, even after the actual process has started. (B) State trajectories and feeding trajectories of off-line optimization and on-line reoptimization. In the presence of PMM, the actual state trajectory resulting from the off-line optimized feeding trajectory deviates from the optimal trajectory (left panels). When the NLMPC repeatedly reoptimizes the feeding trajectory on-line to maximize the final product yield, the deviation of the state trajectory due to PMM can be corrected (right panels).

One of the most significant problems in off-line optimization is the mismatch between predictions made using the models and the actual process states; this is called the process-model mismatch (PMM). When the PMM is large, the off-line-optimized feeding trajectory is no longer optimal in the real process, and the product yield decreases (Fig. 1B).

Mammalian cell culture models generally contain several unknown parameters. When a model-based optimization approach is applied to a process, all unknown parameters in the model must be estimated in advance through experiments using the same cell lines used in the process. Because mammalian cell culture models are generally nonlinear, the design and execution of experiments and parameter estimation should be iterative until the PMM becomes sufficiently small [6,7,9]. If sufficient parameter estimation is unachievable owing to time, economic, or technical reasons, the PMM remains large. In addition, changes in the spatial distribution of metabolite concentrations and cell density in the reactor due to scaling up may also contribute to the PMM. The large PMM attributed to the insufficient parameter estimation and differences between experiments and processes is an obstacle to the application of many excellent models developed by previous studies.

One of the simplest ways to reduce the negative impact of the PMM on feed optimization is to allow the controller to receive real-time measurements of the actual process state and reoptimize the feeding trajectory based on these measurements (Fig. 1A). This approach is called on-line reoptimization, and the controller that solves the optimal control problem in real time is called a nonlinear model predictive controller (NLMPC), which is a type of feedback controller that reflects feedback from the ongoing process in its control inputs.

Several studies have reported the applications of NLMPC in mammalian fed-batch culture [1,10–13]. However, most of them have designed controllers to regulate substrate concentrations [14,15], by-product concentrations [16], substrate consumption rate [17–20], oxygen consumption rate [21], and glycoforms of monoclonal antibodies (mAbs) [22] around fixed set points, instead of maximising the final product yield. Therefore, the resulting feeding trajectories are not always optimal in terms of productivity.

Only a few studies have reported on-line reoptimization of mammalian fed-batch culture with NLMPC designed to maximize product yield. Teixeira et al. demonstrated on-line reoptimization using their hybrid flux balance analysis/neural network model in fed-batch cultures of baby hamster kidney cells [23]. Iyer and Dhir et al. demonstrated on-line reoptimization combined with an on-line parameter estimation algorithm in hybridoma fed-batch culture [24–26]. While these studies showed a certain level of improvement in product yield, it is difficult to distinguish whether it is due to the sophistication of the originally developed model, on-line parameter estimation, or on-line reoptimization alone. Thus, it remains unclear whether on-line reoptimization using a standard kinetic model of mammalian cell culture can successfully compensate for PMM.

This study demonstrated the usefulness and feasibility of a simple on-line reoptimization approach in a scenario where a standard kinetic model is applied to optimize a mammalian fed-batch process after insufficient parameter estimation by simulating two previously reported mAb production processes [2,7]. For each process, we first simulated a single feed case in which the controller optimized the feeding trajectory of a culture medium containing both glucose and glutamine. Next, we extended the simulation to a multiple feed case to demonstrate more complicated control: tthe controller optimized two feeding trajectories, one for a culture medium containing only glucose and the other for a culture medium containing only glutamine. In the simulations, we introduced a PMM by intentionally adding parameter errors to the controller model. Subsequently, we demonstrated the PMM compensation performance of on-line reoptimization by comparing the final product mass with that of off-line optimization in the presence of a PMM. These results confirm that when optimizing mammalian fed-batch culture using a typical kinetic model, on-line reoptimization based on real-time process measurements can improve product yield even if parameter estimation is insufficient.

## Results

We simulated the optimal control using two previously reported mAb production process models: Tremblay’s model [2] and Kontoravdi’s model [7]. The objective of the control was to maximize the mAb mass at the end of the culture period (final mAb mass). We specified the culture conditions and feed constraints of each simulation as the same as those specified in the study that reported the corresponding model. In off-line optimization simulations, the process and controller constitute an open loop. In on-line reoptimization simulations, the process and NLMPC constitute a closed loop (Fig. 1A). The model used to simulate the dynamics of the actual process (the true model) had the parameter values estimated in the original studies. When no PMM was present, the model that the controller/NLMPC used to predict the state trajectories and to optimize the feeding trajectories (controller model) was identical to the true model. When PMM is present, the parameter values of the controller model differ from those of the true model for some parameters (Table 1,2).

### Case 1: Tremblay’s model

Tremblay’s model is a seventh-order model in which both glucose and glutamine concentrations are used to describe the specific growth rate expression. The death rate is governed by the lactate, ammonia, and glutamine concentrations. The specific mAb production rate depends only on the specific growth rate. A detailed description is provided in the Methods and Models section.

Figure 2 shows the process state trajectories predicted by the controller when the feed rate was kept constant in the presence of PMM. The controller predicted an increased production of lactose and ammonia, increased rate of cell death, and decreased production of mAb than that in the actual process.

**Figure 2.**
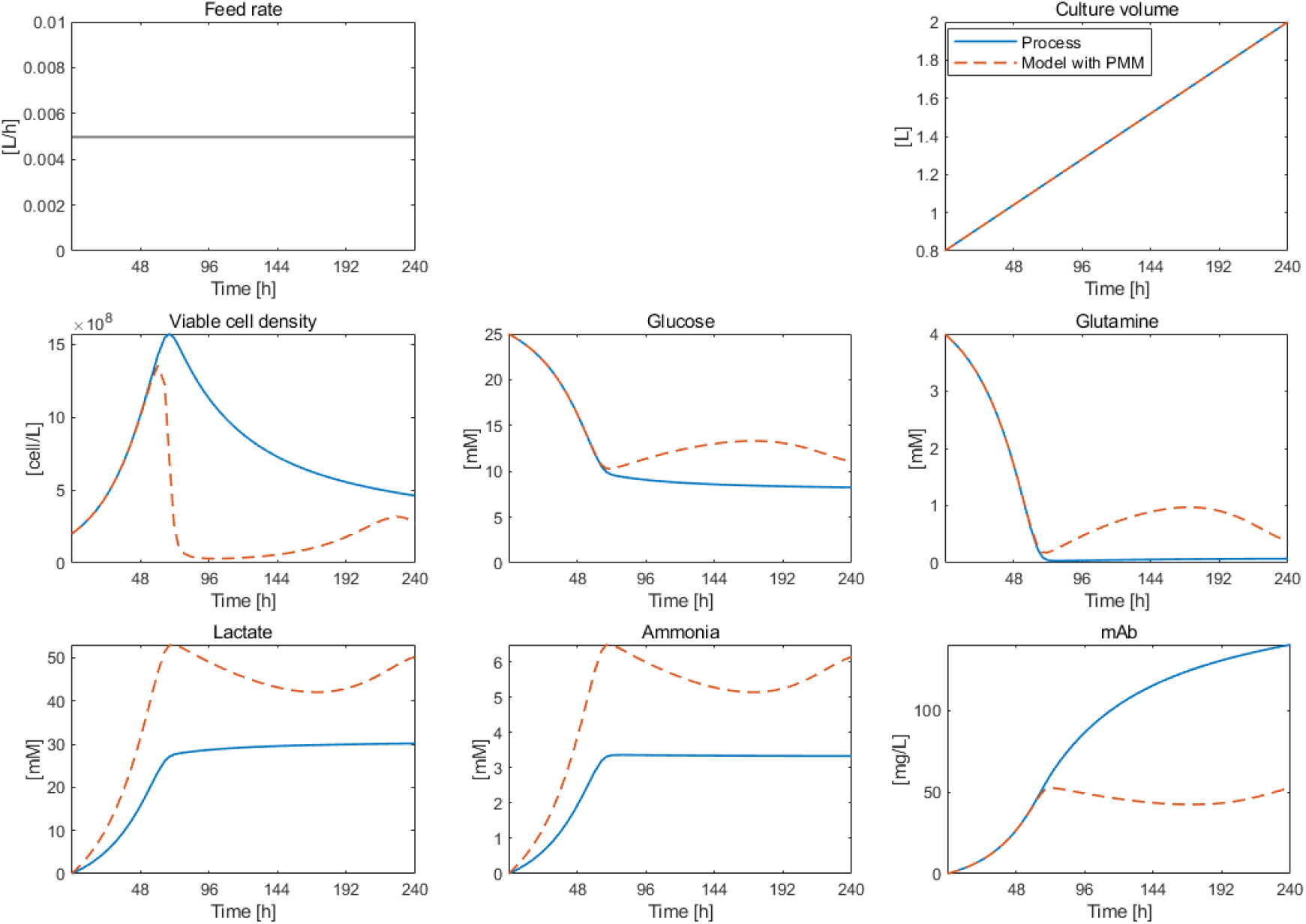
The true (solid lines) and predicted (dotted lines) state variables for the constant feed case in the presence of PMM for Case 1.

#### Case 1-1: Single feed case

The feeding trajectory and resulting state trajectories optimized in the presence or absence of PMM are shown in Figure 3. In the presence of PMM, the off-line controller fed more medium early in the culture than it did in the absence of PMM. Nevertheless, there was no significant difference in the cell growth between the two scenarios. In the presence of PMM, the feed was stopped in the late stage of culture because the culture volume reached the upper limit. This resulted in a faster decrease in the viable cell density at the end of the process, reducing the mAb production rate. When the NLMPC reoptimized the feeding trajectory on-line, the excessive feed early in the culture was suppressed to maintain the feed rate until the end of the process, preventing a faster decrease in viable cell density.

**Figure 3.**
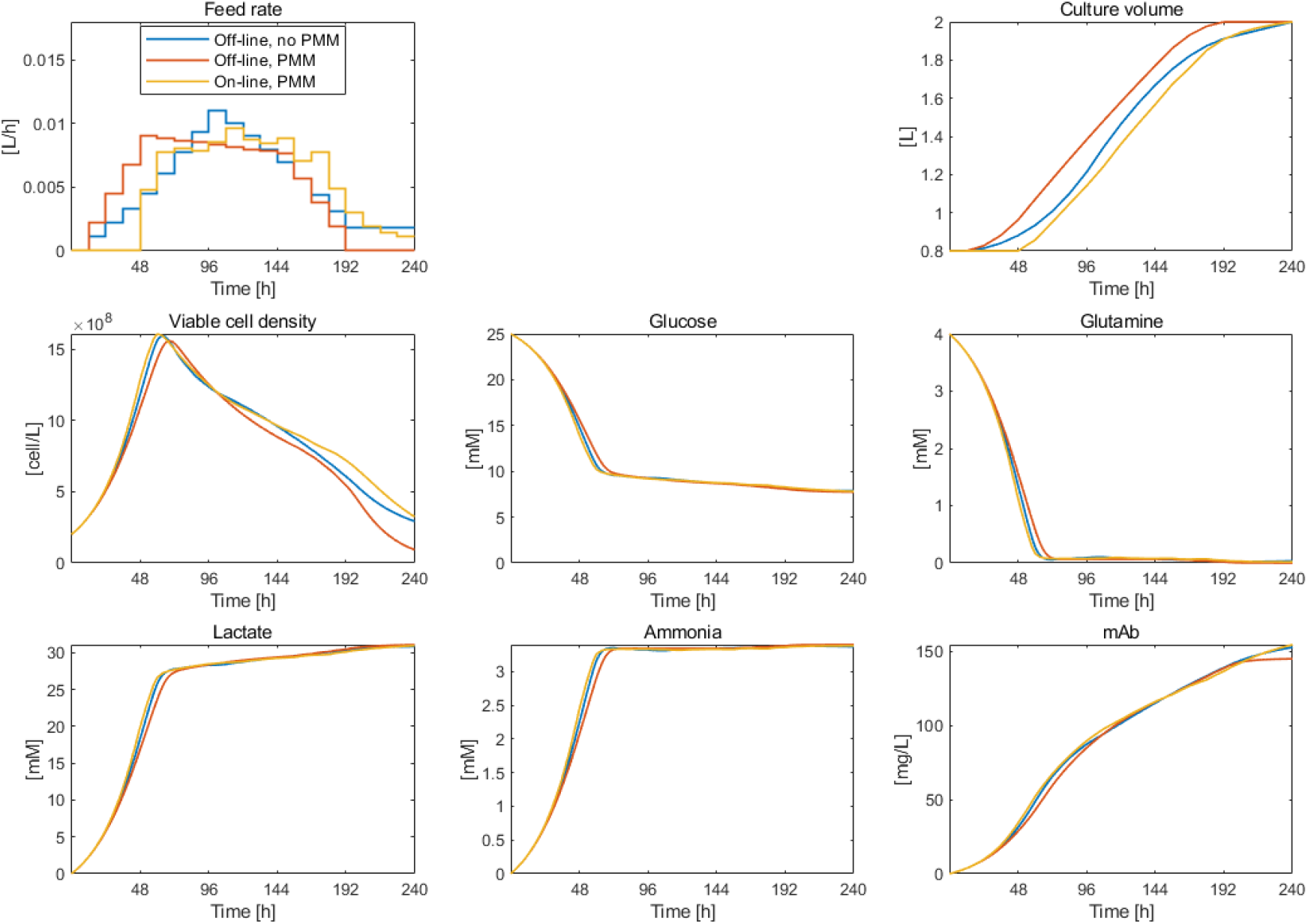
Off-line optimization and on-line reoptimization in the absence or presence of PMM for Case 1-1 (single feed). The blue, red, and yellow lines represent off-line optimization in the absence of PMM, off-line optimization in the presence of PMM, and on-line reoptimization in the presence of PMM, respectively.

Figure 4A shows the final mAb mass in the absence or presence of PMM. In the absence of PMM, off-line optimization increased the final mAb mass by 8.7% compared to the case in which the same volume was fed at a constant rate over the culture duration (constant feed case). In the presence of PMM, off-line optimization was hampered, and the final mAb increase was only 3.1%. When the NLMPC reoptimized the feeding trajectory, the final mAb mass recovered to the same level that was maximized in the absence of PMM, even though the NLMPC could not predict the process state trajectory accurately because of PMM. These results showed that the NLMPC was able to correct the feeding and resulting state trajectories (Fig. 3) and mitigate the reduction in mAb yield due to PMM (Fig. 4A), despite using a model that poorly predicted the process state (Fig. 2).

**Figure 4.**
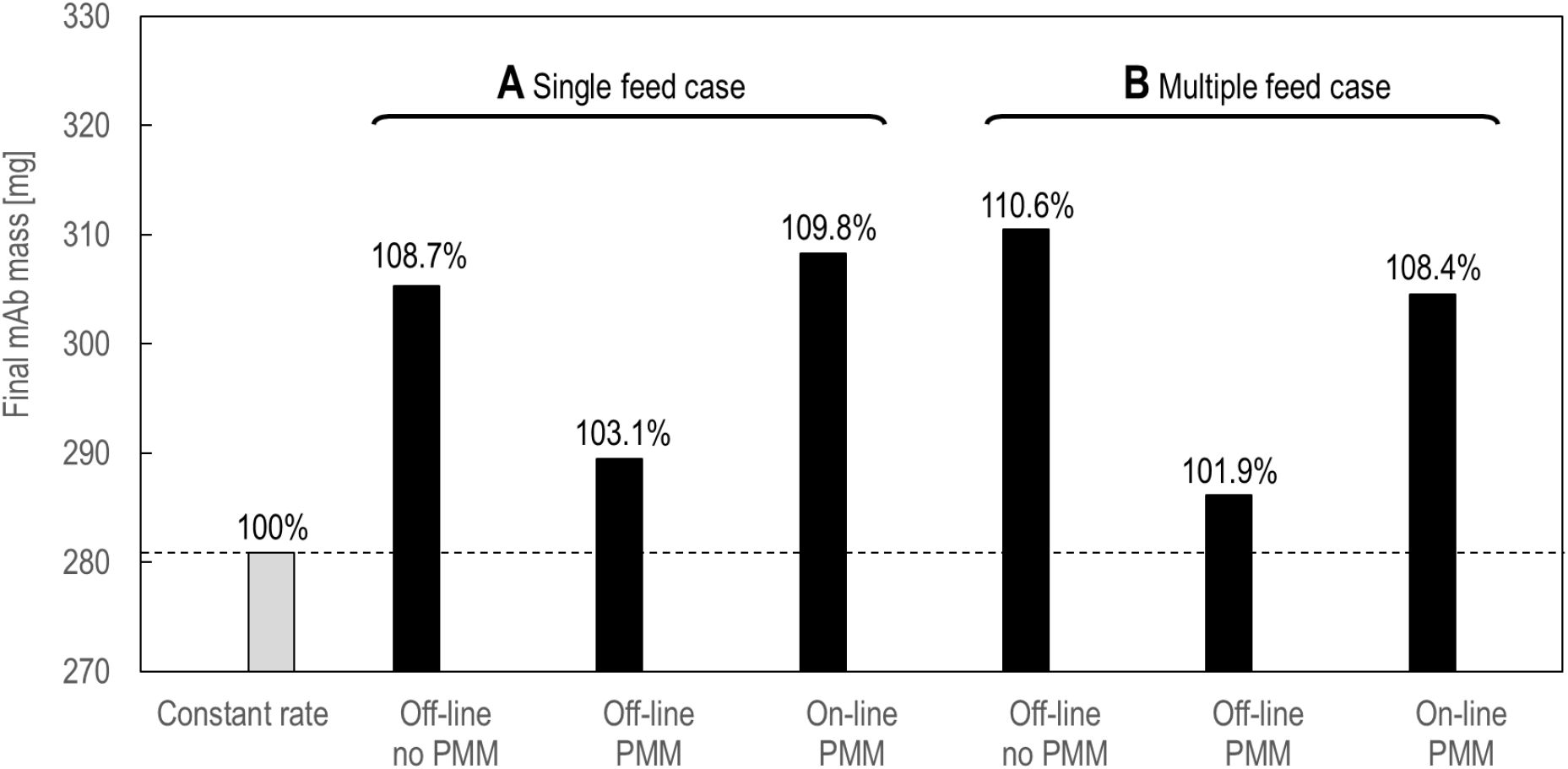
The final mAb mass resulting from off-line optimization and on-line reoptimization for Case 1-1 (A, single feed case) and for Case 1-2 (B, multiple feed case). The grey bar represents the final mAb mass resulting from the constant feed.

#### Case 1-2: Multiple feed case

The feeding trajectory and resulting state trajectories optimized in the presence or absence of PMM are shown in Figure 5. The glucose feeding trajectories off-line-optimized in the presence of PMM were almost the same as those optimized in the absence of PMM. In contrast, in the presence of PMM, glutamine was fed more than in the absence of PMM from 108 to 180 h and was not fed after 192 h. In the on-line reoptimization simulation, as described later in the Methods and Models section, the NLMPC algorithm failed to solve the optimal control problem in two steps (96 h and 108 h) and was not able to update the feed rates. Nevertheless, NLMPC successfully maintained the mAb production rate until the end of the process by suppressing the decrease in the viable cell density.

**Figure 5.**
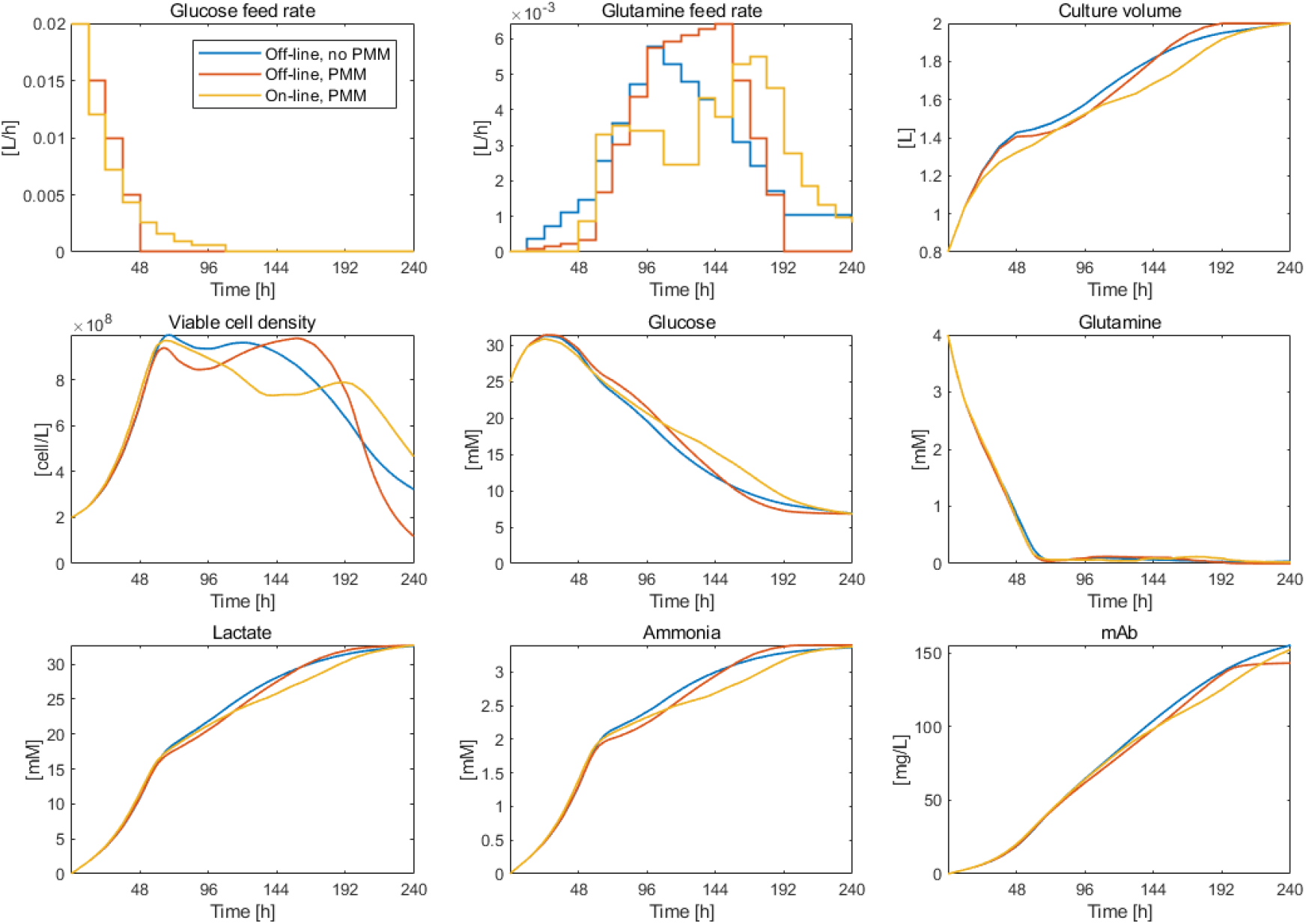
Off-line optimization and on-line reoptimization in the absence or presence of PMM for Case 1-2 (multiple feed case). As in Figure 3, the blue, red, and yellow lines represent off-line optimization in the absence of PMM, off-line optimization in the presence of PMM, and on-line reoptimization in the presence of PMM, respectively.

The final mAb mass was similar to the single feed case: in the absence of PMM, off-line optimization increased the final mAb mass by 10.6% compared to the constant feed case (Fig. 4B). In the presence of PMM, the increase in the final mAb mass was only 1.9%. When the feeding trajectory was reoptimized on-line, as in the single feed case, the final mAb mass recovered to the same level that was maximized in the absence of PMM. These results show that the reoptimization of the feeding trajectory by the NLMPC confirmed in the single feed case can be extended to the multiple-feed case, where more complex control is required.

### Case 2: Kontoravdi’s model

Kontoravdi’s model, as described in detail later in the Methods and Models section, is composed of two main parts: an extracellular submodel and an intracellular submodel. Although the extracellular submodel has the similar structure as the Tremblay’s model, its kinetic rate expressions are different. The intracellular submodel, which outputs the mAb production rate, includes the concentrations of intracellular mRNA and antibody fragments, which are difficult to measure on-line, as the state variables. Therefore, we created a reduced-order model by replacing the intracellular part of the original model with a simple differential equation. There was no significant difference between the state trajectories predicted using the full model and those predicted with the reduced model (Fig. 6). Both models predicted an increased production of lactose and ammonia, increased rate of cell death, and decreased production of mAb than that in the actual process.

**Figure 6.**
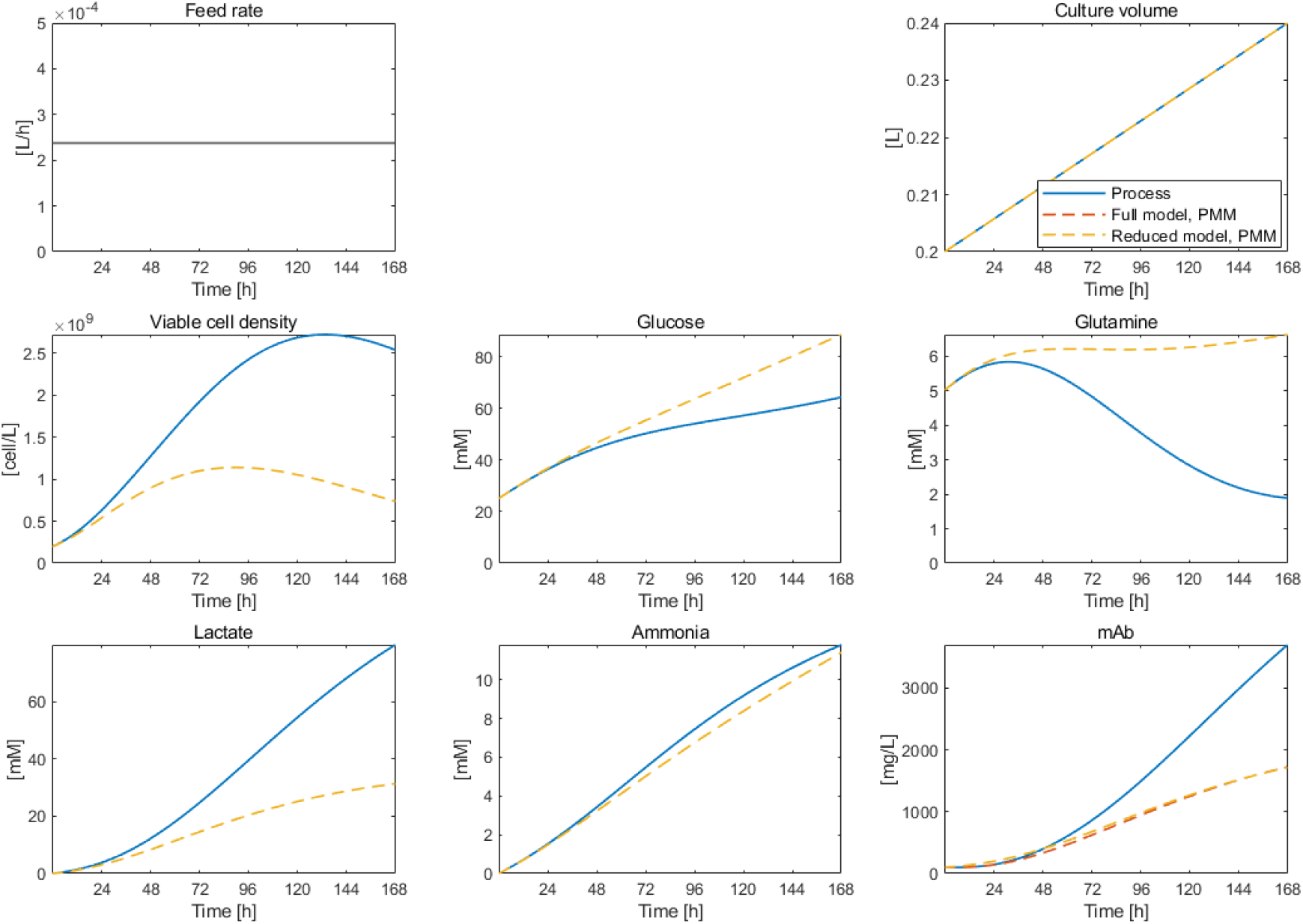
The true (solid lines) and predicted (dotted lines) state variables for the constant feed case in the presence of PMM for Case 2.

#### Case 2-1: Single feed case

In the presence of PMM, the off-line controller decreased the feed rate early in the culture and instead overfed at approximately 120 h (Fig. 7). As a result, cell growth slowed at approximately 72 h. On-line reoptimization successfully corrected underfeeding and overfeeding to prevent the reduction in the cell growth rate.

**Figure 7.**
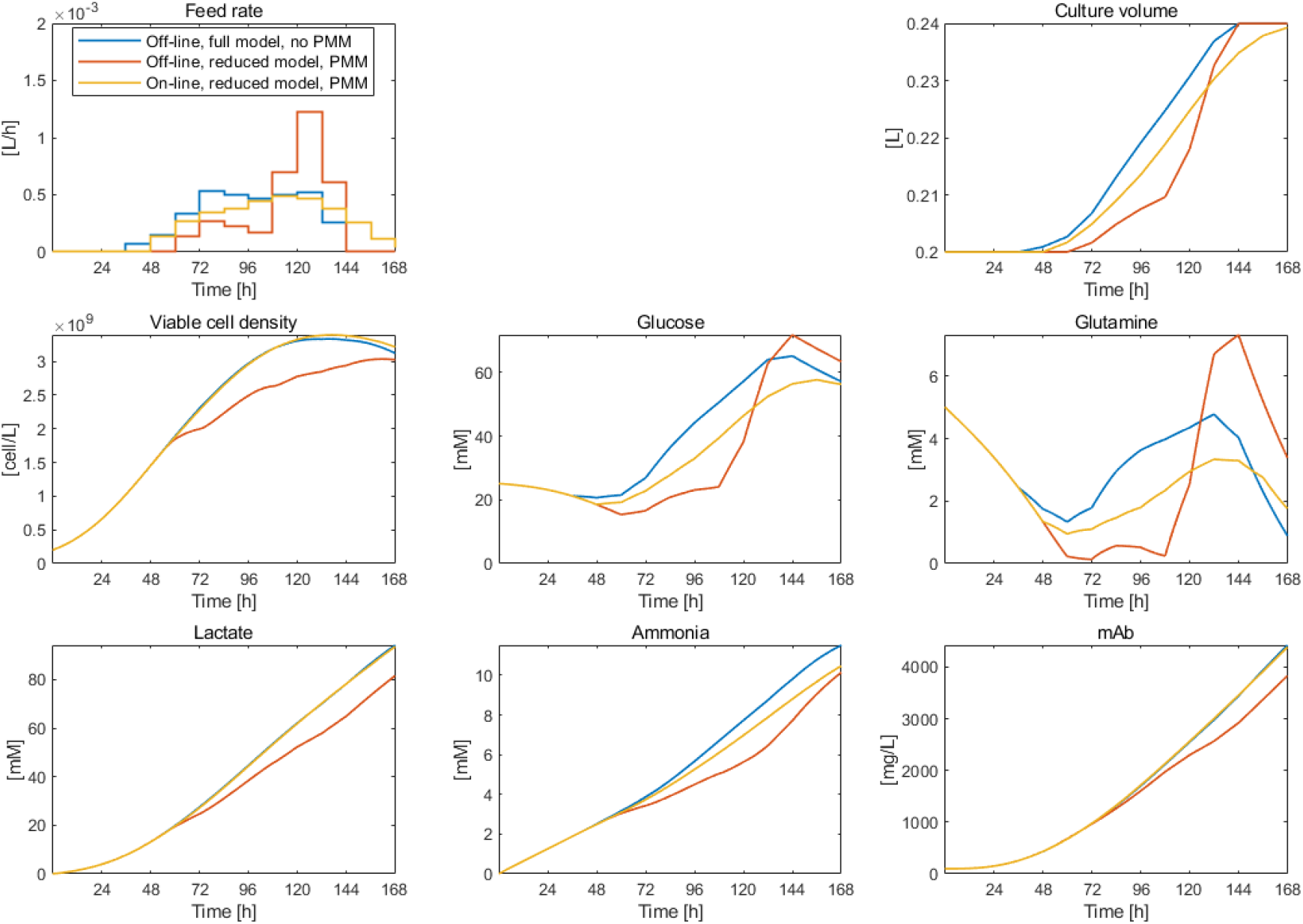
Off-line optimization and on-line reoptimization in the absence or presence of PMM for Case 2-1 (single feed case). As in Figures 3 and 5, the blue, red, and yellow lines represent off-line optimization in the absence of PMM, off-line optimization in the presence of PMM, and on-line reoptimization in the presence of PMM, respectively.

In the absence of PMM, off-line optimization increased the final mAb mass by 19.5% compared to the constant feed case (Fig. 8A). In the presence of PMM, the increase in the final mAb mass decreased to 3.3%. There was little difference in the final mAb mass when the full or reduced model was used for off-line optimization. When the feeding trajectory was reoptimized on-line, as in Case 1-1 and 1-2, the final mAb mass recovered to the same level that was maximized in the absence of PMM.

**Figure 8.**
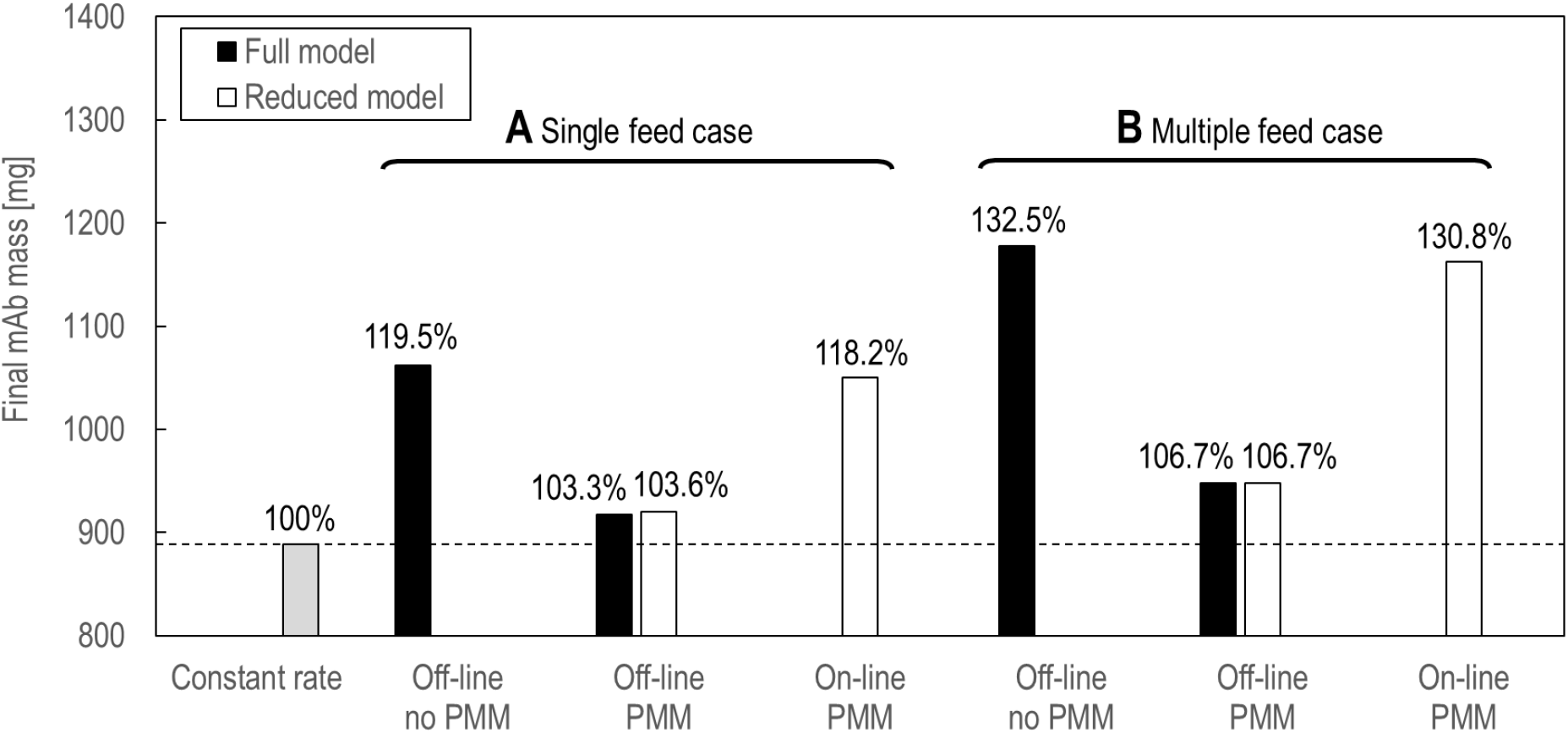
The final mAb mass resulting from off-line optimization and on-line reoptimization for Case 2-1 (A, single feed case) and for Case 2-2 (B, multiple feed case). The grey bar represents the final mAb mass resulting from the constant feed.

#### Case 2-2: Multiple feed case

In the presence of PMM, the glucose feeding trajectory optimized off-line was nearly identical to that optimized in the absence of PMM (Fig. 9). In contrast, glutamine was underfed from 72 to 132 h, which caused a decrease in cell growth. On-line reoptimization successfully corrected glutamine underfeeding and prevented slowdown of cell growth.

**Figure 9.**
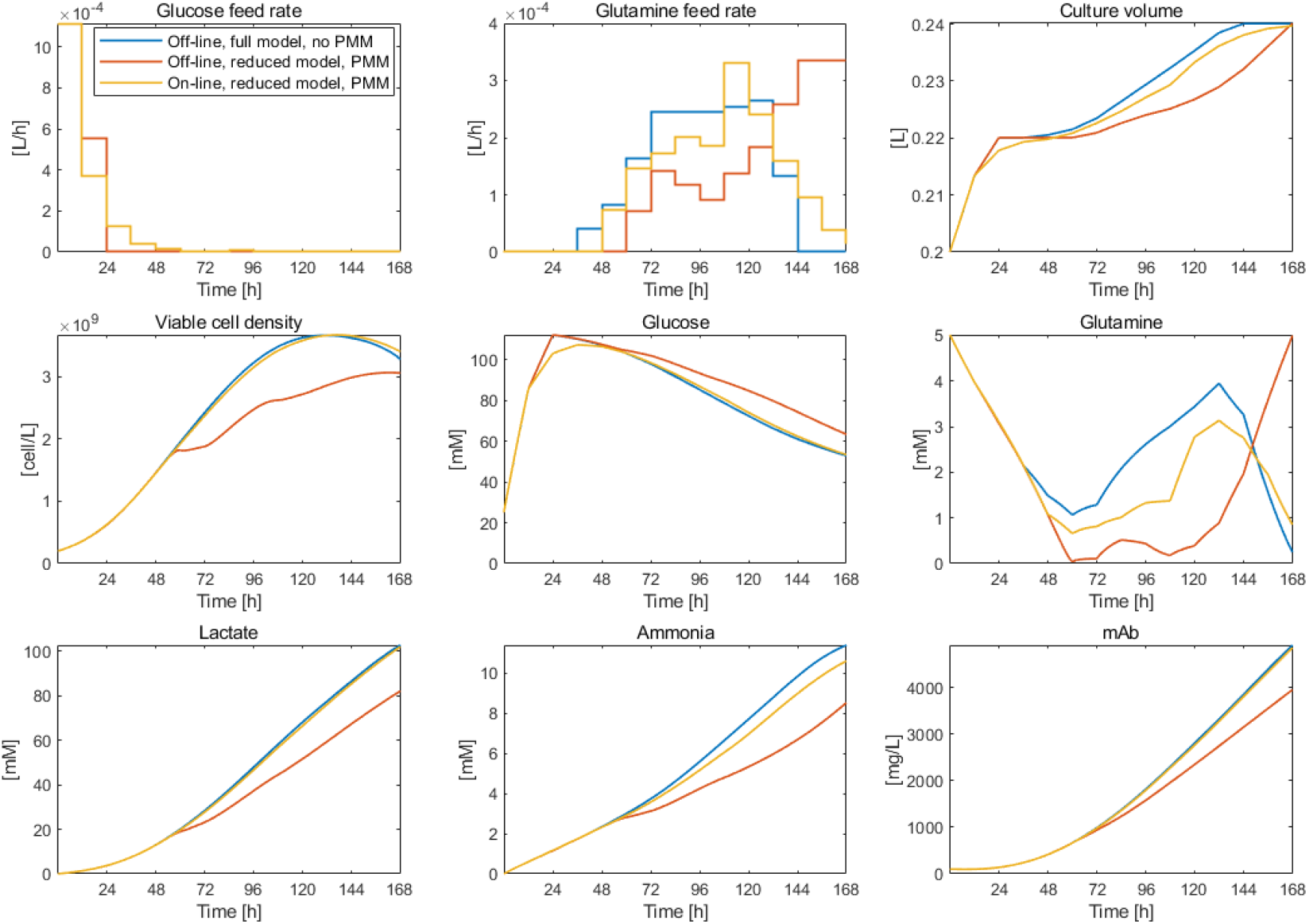
Off-line optimization and on-line reoptimization in the absence and presence of PMM for Case 2-2 (multiple feed case). As in Figures 3, 5, and 7, the blue, red, and yellow lines represent off-line optimization in the absence of PMM, off-line optimization in the presence of PMM, and on-line reoptimization in the presence of PMM, respectively.

In the absence of PMM, off-line optimization increased the final mAb mass even greater than in the single feed case, by 32.5% compared to the constant feed case (Fig. 8). This improvement was largely lost in the presence of PMM. When the feeding trajectories were reoptimized on-line, as in the previous cases, the final mAb mass recovered to the same level that was maximized in the absence of PMM. From these results, we can expect that the on-line reoptimization approach is applicable not only to Tremblay’s model but also to other typical kinetic models.

## Discussion

In this study, we verified the usefulness and feasibility of an on-line reoptimization approach using NLMPC in mammalian fed-batch culture. In this approach, the NLMPC repeatedly solves the optimal control problem formulated to maximize the final product mass based on the measured values of the state variables as initial states at each sample time. This approach was demonstrated through simulations using two previously reported mAb production process models. In both models, our approach successfully suppressed the loss of mAb yield due to the intentional introduction of the PMM.

Both models used in this study are relatively simple mammalian models developed to date. When using more complex models, the computational burden is a potential problem. However, in this study, the optimal control problem was solved using a standard personal computer with an Intel Core i5 processor and 8GB DDR4 SDRAM in considerably less time than the 12-hour sample time (approximately ten seconds for the single feed cases and approximately one minute for the multiple feed cases). Thus, there is still an ample margin for computation time. Furthermore, if parallel computing with a multinode computer cluster is available, the optimal control problem can be solved in a shorter time. Therefore, we believe that the simple approach of on-line reoptimization can be applied to even more complex models such as those involving glycosylation [27–34].

In the simulations, the intentionally introduced PMM was sufficiently large that the state trajectories predicted by the controllers deviated significantly from the true trajectories (Figs 2 and 6). In particular, the predicted final mAb concentration for the constant feed rate was less than half of the true value in both cases 1 and 2. As a result, off-line optimization was significantly disturbed such that the final mAb mass dropped to the same level as that for the constant feed case (Fig. 4 and 8). Nevertheless, the proposed approach could cope with such a large PMM.

In Case 1-1, the mAb yield resulting from off-line optimization in the absence of PMM was slightly lower than that resulting from on-line reoptimization. This is likely due to constraints on the feeding trajectory, as described in the Methods and Models section. To improve the robustness of the solver, we reduced the decision frequency of the feed rate from every step to every two steps. The feed rates of the in-between steps were set to the average of the feed rates of the previous and subsequent steps (linear interpolation). As a result, the degree of freedom in the feeding trajectories was reduced, along with the final mAb mass. In on-line reoptimization, the feed rate was independently determined for each step, and thus, the mAb mass can exceed the final mAb mass maximized in the absence of PMM.

In the simulations, the NLMPC used viable cell density and glucose, glutamine, lactose, and ammonia concentrations measured every 12 h. In real processes, these variables can be measured within a few minutes using commercially available automated cell culture analysers such as BioProfile FLEX2 (Nova Biomedical). Thus, the proposed approach can easily be applied to real bioprocesses. Some mechanistic models include state variables that are difficult to measure on-line, such as the amount of mRNA and protein in cells. We believe that this approach can also be applied to such models by reducing the order of the models, as we did for Kontoravdi’s model, or by combining NLMPC with state estimation algorithms such as the Kalman filter and particle filter.

## Methods and Models

### Optimal control problem

We designed an optimal control problem such that the mAb mass at the end of the culture was maximized. The upper limit of the feed volume was set as an optimization constraint. Thus, the optimal control problem was formulated for the single feed case as:

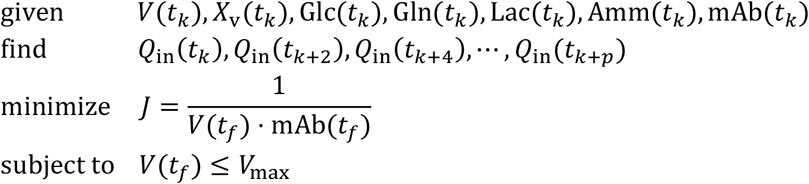

and for the multiple feed case as:

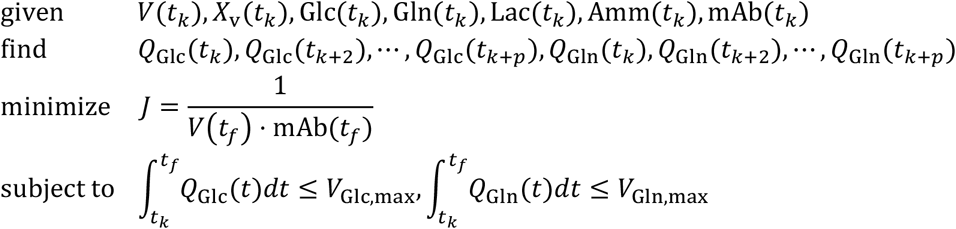

where *k* is the current step, *t_k_* is the current sample time, *t_f_* is the culture duration, *t_s_* is the sampling interval, and 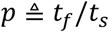 is the prediction horizon. The sampling interval was set to 12 h. To reduce the computational load, we decimated the decision variables. The feed rate values for every two steps (24 h) were taken as decision variables, and the feed rate values at the in-between steps were linearly interpolated.

We used the Nonlinear MPC object *nlmpc* in Model Predictive Control Toolbox of MATLAB/Simulink for the simulations. The *nlmpc* object requires the model to be discretised at the sample time. Therefore, we divided the sample time into 100 substeps and numerically integrated the model using the forward Euler method. To solve the optimal control problem, we used the default solver of *nlmpc*. This solver is a so-called direct method that transforms the optimal control problem into a nonlinear programming problem constrained by the state equations (model) with the state variables and input variables (feed rates) at each sample time as decision variables. The solver was set to use the sequential quadratic programming (SQP) algorithm to solve the nonlinear programming problem. The solver may fail to find a feasible solution depending on the measured values of the state variables at certain sample times. At these times, the NLMPC was set to continue feeding at the same rate as in the previous step. To improve the robustness of the solver, we set the feeding trajectory and state trajectory computed in the previous step as the initial guess for the next step, which is called the warm start.

### Case 1: Tremblay’s model

#### Model

Tremblay’s model contains seven state variables. The state variables and model parameters are listed in Tables 1 and 2, respectively. The controller model had different parameter values from those of the model used to simulate the process (true model).

The ordinary differential equations for the single feed case are:

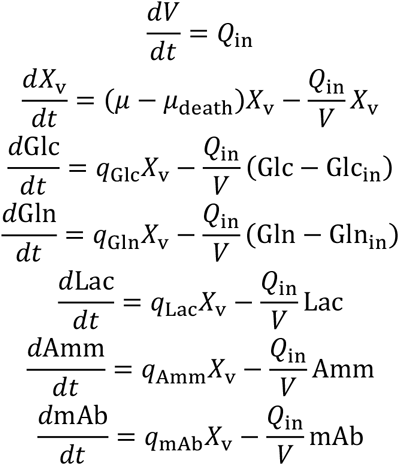

The ordinary differential equations for the multiple feed case are:

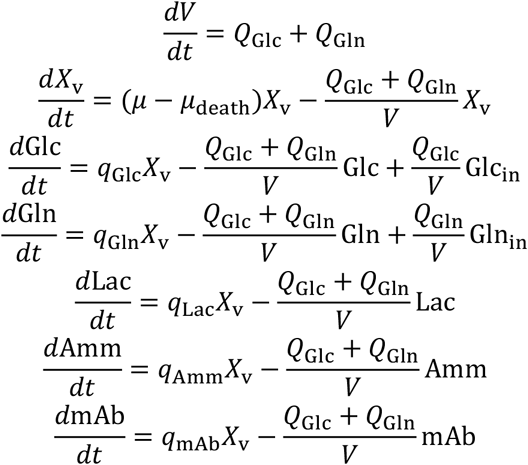

The cell growth and death rates are:

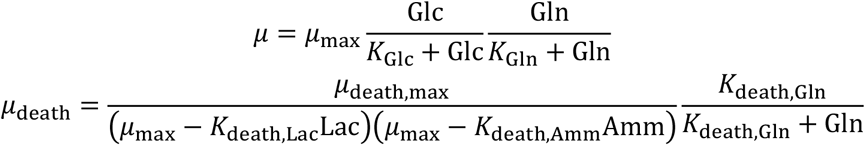

The metabolite production rates are:

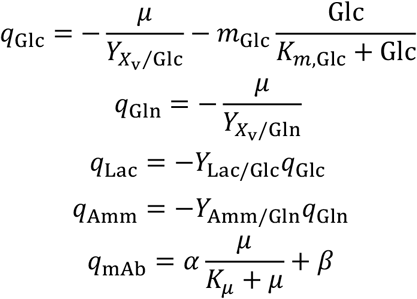

The negative sign represents consumption.

#### Culture conditions

The culture duration is 240 h. The initial culture conditions are:

- V0 = 0.8 L
- Xv0 = 2.0E+8 cell/mL
- Glc0 = 25 mM
- Gln0 = 4 mM
- Lac0 = 0 mM
- Amm0 = 0 mM
- mAb0 = 0 mg/L

For the single feed case, the feed concentrations of glucose and glutamine are:

- Glc_in = 25 mM
- Gln_in = 4 mM

The maximum feed volume is 1.2 L.

For the multiple feed case, the feed concentrations of glucose and glutamine are:

- Glc_in = 50 mM
- Gln_in = 8 mM

The maximum feed volume is 0.6 L for both the glucose and glutamine feed.

### Case 2: Kontoravdi’s model

#### Model

Kontoravdi’s model consists of an extracellular submodel and an intracellular submodel. The state variables are listed in Tables 1 and 3, and the model parameters are listed in Table 4. The controller model has different parameter values from the model used for the process simulation (true model).

#### Extracellular submodel

The extracellular submodel contains the same state variables as Tremblay’s model does (Table 1). The ordinary differential equations for the single feed case are:

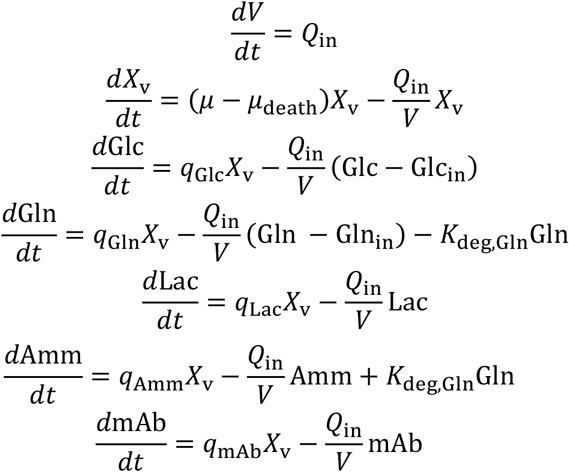

The ordinary differential equations for the multiple feed case are:

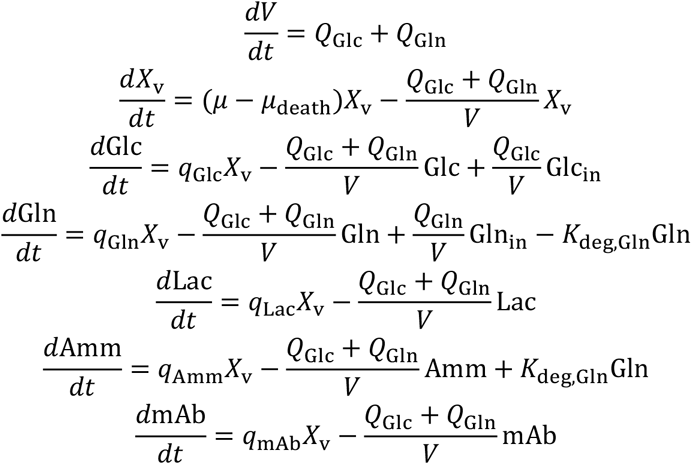

The cell growth and death rates are:

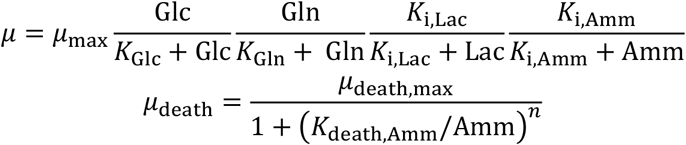

The metabolite production rates are:

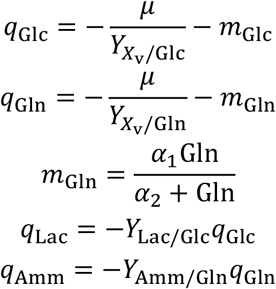

The negative sign represents consumption.

#### Intracellular submodel

The intracellular submodel contains eight state variables (Table 3). In this submodel, genes encoding the light and heavy chains of mAb are transcribed and then translated. The light and heavy chains are assembled into mAb molecules and secreted from the cell via the Golgi apparatus. Figure S2 illustrates the reaction network of the submodel.

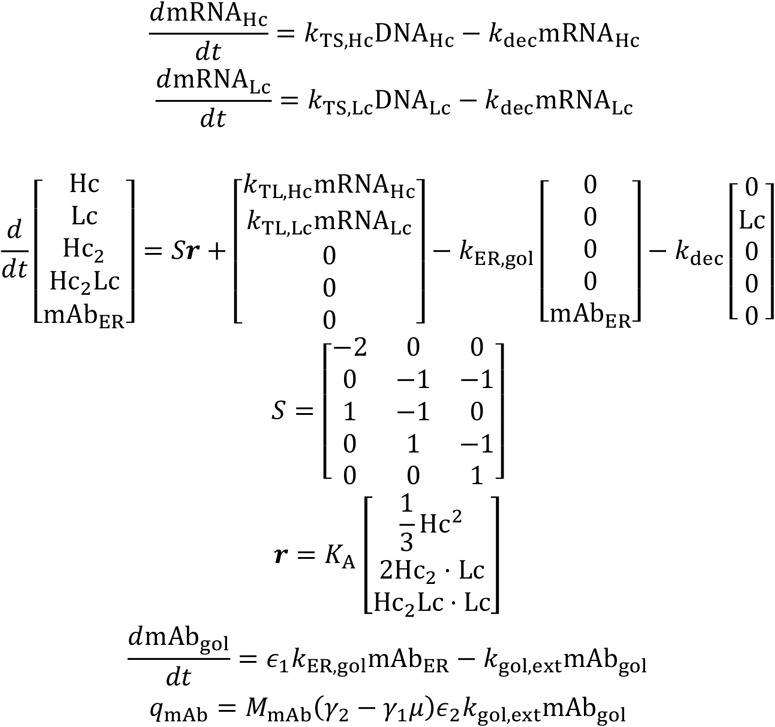

#### Model reduction

The intracellular submodel defines the intracellular amounts of mRNAs and mAb fragments as state variables. To perform on-line reoptimization, all state-variable measurements must be passed to the NLMPC at each sample time.

However, it is difficult to measure the concentration of these intracellular mRNAs and proteins every 12 h in the real process. Therefore, we replaced the submodel with a simple differential equation to reduce the model, and used the reduced model as a controller model of the NLMPC. The differential equation was adopted from the Chinese hamster ovary cell culture model reported previously [27].

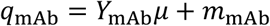

The parameter values were estimated using Simulink Design Optimization to minimize the sum of the squared errors (SSE) between the predictions of mAb concentration by the full model and the reduced model. The SSE was calculated from five simulations for the constant feed case with different feed rates (no feed, 2.98E-5, 6.00E-5, 1.19E-4, and 2.38E-4 L/h). We used a nonlinear least-squares solver *lsqnonlin* as the optimization solver. The estimated parameter values are:

- Y_mAb = 3.0583E-12
- m_mAb = 1.1989E-08

#### Culture conditions

The culture duration is 168 h. The initial culture conditions are:

- V0 = 0.20 L
- Xv0 = 2.0E+8 cell/mL
- Glc0 = 25.1 mM
- Gln0 = 5.01 mM
- Lac0 = 0 mM
- Amm0 = 0 mM
- mAb0 = 100 mg/L

For the single feed case, the feed concentrations of glucose and glutamine are:

- Glc_in = 500 mM
- Gln_in = 100 mM

The maximum feed volume is 0.04 L.

For the multiple feed case, the feed concentrations of glucose and glutamine are:

- Glc_in = 1000 mM
- Gln_in = 200 mM

The maximum feed volume is 0.02 L for both the glucose and glutamine feed.

## Code availability

Simulation code can be downloaded the following URL on GitHub (https://github.com/kkunida/202212_Ohkubo_bioRxiv.git).

## Acknowledgments

We would like to thank Dr. Masaaki Nagahara from the University of Kitakyushu for helpful and technical comments. We thank Editage (www.editage.com) for English language editing. This study was supported by the Japan Society for the Promotion of Science (JSPS) KAKENHI Grant Numbers 19K20400 (K.K.) and Next Generation Interdisciplinary Research Project of Nara Institute of Science and Technology (NAIST).

## Author contributions

T. O. and K. K. designed the project; T.O. performed the modeling and simulation; T.O., Y. S., and K. K. prepared the manuscript.

