## Supplemental Figures and Tables for "On-line reoptimization of mammalian fed-batch culture using a nonlinear model predictive controller"

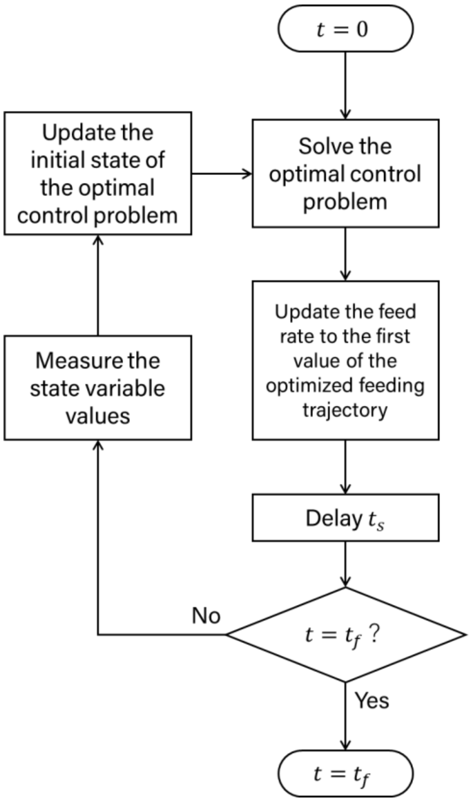

Figure S1. The flow chart of on-line reoptimization.

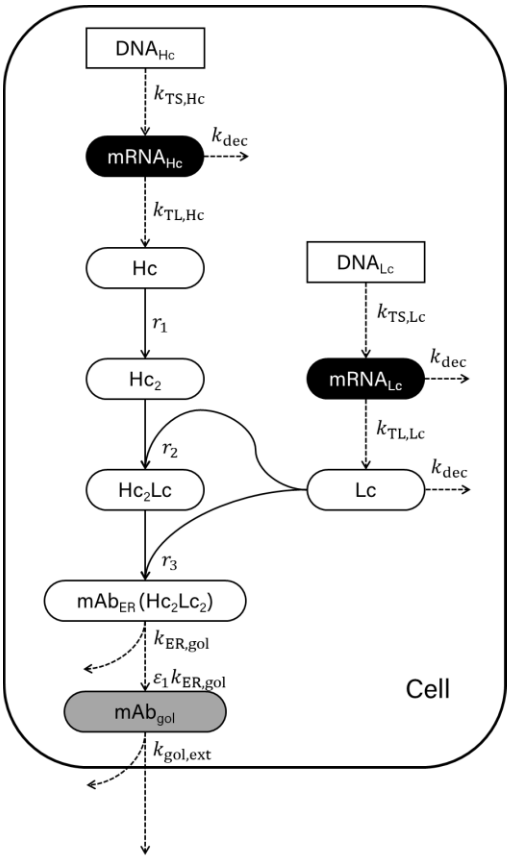

$$q_{\mathrm{mAb}} = M_{\mathrm{mAb}}(\gamma_2 - \gamma_1 \mu)\varepsilon_2 k_{\mathrm{gol,ext}} \mathrm{mAb}_{\mathrm{gol}}$$

Figure S2. The reaction network of the intracellular submodel of Kontoravdi's model.

Table 1. Notation and description of the state variables in Tremblay's model (Case 1), and the extracellular submodel of Kontoravdi's model (Case 2).

| Notation | Unit | Description |
|---|---|---|
| $V$ | L | Volume of culture |
| $X_{\mathrm{v}}$ | cell/L | Viable cell density |
| $\mathrm{Glc}$ | mM | Glucose concentration |
| $\mathrm{Gln}$ | mM | Glutamine concentration |
| $\mathrm{Lac}$ | mM | Lactate concentration |
| $\mathrm{Amm}$ | mM | Ammonia concentration |
| $\mathrm{mAb}$ | mg/L | mAb concentration |

Table 2. The model parameters of Tremblay's model (Case 1). The parameter values used to simulate the process (true parameters) were those estimated in the original study. In the controller model, four parameters were set differently to introduce PMM (blank represents the same value as the process).

| Notation | Unit | Process (true parameters) | Controller model |
|---|---|---|---|
| $\mu_{\mathrm{max}}$ | /h | 4.54E-2 | |
| $\mu_{\mathrm{death,max}}$ | /h^3 | 4.99E-05 | |
| $K_{\mathrm{Glc}}$ | mM | 1 | |
| $K_{\mathrm{Gln}}$ | mM | 0.3 | |
| $K_{\mathrm{death,Lac}}$ | /(mM*h) | 4.17E-4 | 8.33E-4 |
| $K_{\mathrm{death,Amm}}$ | /(mM*h) | 2.5E-3 | 5E-3 |
| $K_{\mathrm{death,Gln}}$ | mM | 0.02 | |
| $Y_{X_{\mathrm{v}}/\mathrm{Glc}}$ | cell/mmol | 1.09E+8 | |
| $Y_{X_{\mathrm{v}}/\mathrm{Gln}}$ | cell/mmol | 3.8E+8 | |
| $Y_{\mathrm{Lac/Glc}}$ | mmol/mmol | 1.8 | 3.6 |
| $Y_{\mathrm{Amm/Gln}}$ | mmol/mmol | 0.85 | 1.7 |
| $m_{\mathrm{Glc}}$ | mmol/(cell*h) | 7.08E-11 | |
| $K_{m,\mathrm{Glc}}$ | mM | 19 | |
| $K_{\mu}$ | /h | 8.33E-4 | |
| $\alpha$ | mg/(cell*h) | 1.07E-9 | |
| $\beta$ | mg/(cell*h) | 1.46E-10 | |

Table 3. Notation and description of the state variables in the intracellular submodel of Kontoravdi's model (Case 2).

| Notation | Unit | Description |
|---|---|---|
| $\mathrm{mRNA_{Hc}}$ | molecule/cell | Number of mRNA molecules encoding the heavy chain |
| $\mathrm{mRNA_{Lc}}$ | molecule/cell | Number of mRNA molecules encoding the light chain |
| $\mathrm{Hc}$ | molecule/cell | Number of the heavy chain molecules |
| $\mathrm{Lc}$ | molecule/cell | Number of the light chain molecules |
| $\mathrm{Hc_2}$ | molecule/cell | Number of the heavy chain dimers |
| $\mathrm{Hc_2Lc}$ | molecule/cell | Number of the heavy and light chain complexes |
| $\mathrm{mAb_{ER}}$ | molecule/cell | Number of mAb molecules in the endoplasmic reticulum |
| $\mathrm{mAb_{gol}}$ | molecule/cell | Number of mAb molecules in the Golgi apparatus |

Table 4. The model parameters for Case 2 (Kontoravdi's model). The parameter values used to simulate the process (true parameters) were those estimated in the original study. In the controller model, four of the parameters were set differently to introduce PMM (blank represents the same value as the process).

| Notation | Unit | Process (true parameters) | Controller model |
|---|---|---|---|
| Extracellular | | | |
| $\mu_{\mathrm{max}}$ | /h | 0.058 | 0.0522 |
| $\mu_{\mathrm{death,max}}$ | /h | 0.03 | |
| $K_{\mathrm{Glc}}$ | mM | 0.75 | |
| $K_{\mathrm{Gln}}$ | mM | 0.075 | |
| $K_{\mathrm{i,Lac}}$ | mM | 171.76 | 85.88 |
| $K_{\mathrm{i,Amm}}$ | mM | 28.48 | 14.24 |
| $K_{\mathrm{death,Amm}}$ | mM | 1.76 | |
| $n$ | dimensionless | 2 | |
| $Y_{X_{\mathrm{v}}/\mathrm{Glc}}$ | cell/mmol | 2.6E+8 | |
| $Y_{X_{\mathrm{v}}/\mathrm{Gln}}$ | cell/mmol | 8E+8 | |
| $m_{\mathrm{Glc}}$ | mmol/(cell*h) | 4.9E-14 | 9.8E-14 |
| $\alpha_1$ | mmol/(cell*h) | 3.4E-13 | |
| $\alpha_2$ | mM | 4 | |
| $Y_{\mathrm{Lac/Glc}}$ | mmol/mmol | 2 | |
| $Y_{\mathrm{Amm/Gln}}$ | mmol/mmol | 0.45 | |
| $K_{\mathrm{deg,Gln}}$ | mM | 9.6E-3 | |
| Intracellular | | | |
| $k_{\mathrm{TS,Hc}}$ | molecule/(molecule*h) | 3000 | |
| $k_{\mathrm{TS,Lc}}$ | molecule/(molecule*h) | 4500 | |
| $\mathrm{DNA_{Hc}}$ | molecule/cell | 100 | |
| $\mathrm{DNA_{Lc}}$ | molecule/cell | 100 | |
| $k_{\mathrm{dec}}$ | /h | 0.1 | |
| $k_{\mathrm{TL,Hc}}$ | molecule/(molecule*h) | 17 | |
| $k_{\mathrm{TL,Lc}}$ | molecule/( molecule*h) | 11.5 | |
| $K_{\mathrm{A}}$ | cell/(molecule*h) | 1E-6 | |
| $k_{\mathrm{ER,gol}}$ | /h | 0.69 | |
| $k_{\mathrm{gol,ext}}$ | /h | 0.14 | |
| $\epsilon_1$ | dimensionless | 0.995 | |
| $\epsilon_2$ | dimensionless | 1 | |

| | | |
|---|---|---|
| $\gamma_1$ | h | 0.1 |
| $\gamma_2$ | dimensionless | 1 |
| $M_{\mathrm{mAb}}$ | mg/molecule | 2.4908084E-16 |
